# An altered metabolism could contribute to the low activation of neonatal CD8^+^T cells

**DOI:** 10.1101/726885

**Authors:** Sánchez-Villanueva José Antonio, Rodríguez-Jorge Otoniel, Ramírez-Pliego Oscar, Rosas Salgado Gabriela, Abou-Jaoudé Wassim, Hernandez Céline, Naldi Aurélien, Thieffry Denis, Santana María Angélica

**Author notes:** **Corresponding authors:** Santana M.A. < >, Thieffry D. < >.

## Abstract

A low response of neonatal T cells to infections contributes to a high incidence of morbidity and mortality of neonates. Here we evaluated the impact of the cytoplasmic and mitochondrial levels of Reactive Oxygen Species (ROS) of adult and neonatal CD8^+^T cells on their activation potential. We further constructed a logical model to decipher the interplay of metabolism and ROS on T cell signaling. Our model captures the interplay between antigen recognition, ROS and metabolic status in T cell responses. This model displays alternative stable states corresponding to different cell fates, i.e. quiescent, activated and anergic states, depending on ROS status. Stochastic simulations with this model further indicate that differences in ROS status at the cell population level tentatively explain the lower activation rate of neonatal compared to adult CD8^+^T cells upon TCR engagement.

## Introduction

Infections in children under six months cause around four million deaths per year (PrabhuDas et al., 2011). Neonatal T cells have low, tolerant or skewed response and a relatively high threshold of activation, potentially involving epigenetic mechanisms (Adkins, 2005; Fike et al., 2019; Marchant and Goldman, 2005; Ndure and Flanagan, 2014). Molecules present in the neonatal serum contribute to the low response of neonatal cells, among them adenosine (Hasko and Cronstein, 2004; Levy et al., 2006), arginase (Elahi, 2014), and other proteins (Chatterton et al., 2013; Levy, 2007). Arginase and adenosine are metabolic inhibitors associated with T cell un-responsiveness in cancer (Liu et al., 2010; Monticelli et al., 2016; Vaupel and Multhoff, 2016; Wang et al., 2014). CD8^+^T cells from newborns have distinct transcriptional and epigenetic profiles biased towards innate immunity and, albeit in homeostatic proliferation, their clonal expansion and effector functions are diminished (Galindo-Albarran et al., 2016). Lower mitochondria mass and membrane potential, as well as differences in calcium fluxes and in calcium and potassium channels have been reported (Meszaros et al., 2015; Palin et al., 2013; Toldi et al., 2010). Naïve, memory and effector functions depend on different metabolic programs, adapted to cellular requirements. Naïve T cells are metabolically dormant (quiescent). After antigen encounter, they may either tolerate the antigen, become activated, or turn anergic. Despite important advances in the field of immunometabolism, a comprehensive view of the interplay between antigen recognition, metabolic status and ROS is missing. Our study combines experimental measurements with computational modeling to assess the role of metabolism, in particular of ROS, on T cell signaling.

We evaluated cytosolic (c) and mitochondrial (m) ROS levels on naïve CD8^+^T cells from human newborns and adults, at basal level and after TCR stimulation. Our results suggest fundamental differences in the ROS signaling and redox status between CD8^+^T cells from newborns and adults. Alterations of Redox and metabolic nodes in the model result in low neonatal CD8^+^T cells response, establishing the involvement of the metabolic status of the neonatal cells in their impaired response.

Using a logical formalism implemented in the software GINsim ((Naldi et al., 2018a) http://ginsim.org), we defined a comprehensive dynamical model integrating the most relevant signaling, metabolic and transcriptional regulatory components controlling the activation of CD8^+^T lymphocytes.

This model displays alternative stable states, which corresponds to different cell fates, i.e. quiescence, activation and anergy, depending on ROS status. Stochastic simulations further suggest that the lower activation rate of neonatal compared to adult CD8^+^T cells upon TCR engagement is attributable to differences in ROS status at the cell population level.

## Materials and methods

### Blood collection and cell purification

Cord blood was collected at Hospital General de Cuernavaca Dr. José G. Parres, with informed mothers’ consent. Adult samples were obtained from leukocyte concentrates at the Centro Estatal de Transfusión Sanguínea. Samples were immediately processed. CD8^+^T cells were purified as previously described (Hernandez-Acevedo et al., 2019). Briefly, the blood was centrifuged on Lymphoprep™ (Axis-Shield, UK) and Mononuclear cells were incubated with 1 ml of erythrocytes and the RossetteSep™ CD8^+^T cell enrichment cocktail to obtain Total CD8^+^T cells (15063; StemCell Technologies, Canada). The memory cells were eliminated using magnetic beads (8803; Pierce; Thermo Fisher Scientific, Bremen, Germany) loaded with a CD45RO-specific mAb (eBiosciences, San Diego, CA). We obtained ≥94% naïve CD8^+^T cells.

### Cell stimulation

Naïve CD8^+^T cells were cultured in RPMI containing 1% L-glutamine, antibiotics (100 U/mL penicillin and 100 μg/mL streptomycin) and 5% fetal calf serum. For stimulation 1 μg/mL or each, anti-CD3/anti-CD28 mAbs was used (OKT3, 70-0037-U100, CD28.2, 70-0289-U100, Tonbo Biosciences), cross-linked with anti-mouse mAb (405301, BioLegend). Cells were incubated for 6 h under 5% CO_2_ at 37°C.

### Flow cytometry

For activation assessment the anti-CD69 FITC (Genetex GTX43516) was used. Cells were stained as previously described [15].

For Redox measurements, cells were incubated for 15 minutes at 37°C in staining buffer with either: 2.5 μM of MitoSOX Red Mitochondrial Superoxide Indicator (M36008 ThermoFisher Scientific); or 2 μM of Dihydroethidium (DHE) (D11347 ThermoFisher Scientific); or 5 μM of Carboxy-H_2_DCFDA (C400 ThermoFisher Scientific). Cells were washed twice with staining buffer.

Flow cytometry was assessed on a FACScalibur cytometer using the CellQuest software. The software FlowJo (Tree Star, CA) was used for analysis.

### Statistical analysis

An unpaired Mann-Whitney test was used for comparison between samples, or the Wilcoxon test for comparison between paired samples.

### Logical modeling

To model the T cell network, we first defined a *regulatory graph*, where each node represents a component of the signaling network. Most nodes are associated with Boolean variables, but some are allowed to take three levels of activity (0, 1 and 2) when biologically justified. Nodes are connected through arcs, which represent the regulatory influences between them, positive or negative. Next, we defined logical rules to determine the level of activity of each target node as a function of the levels of its regulators (Suppl. Table 1).

We used the GINsim software (v3.0.0b, [19], http://ginsim.org) to build the TCR-REDOX-Metabolism signaling model and to identify its stable states.

We used the MaBoSS software (v2.0, (Stoll et al., 2012)) to perform stochastic simulations of our logical model. These simulations use default parameters: 50’000 runs with identical up and down rates for all components.

All our model analyses are reproducible using an interactive Python notebook and virtual software environment, available together with the model file on GINsim repository (http://ginsim.org/node/229) (Naldi et al., 2018b).

## Results

### Evaluation of ROS in neonatal and adult CD8+T cells

We purified naïve CD8^+^T cells from neonate and adult donors as previously described (Hernandez-Acevedo et al., 2019). Glycolysis enzymes are over-expressed in the neonatal CD8^+^T cells, which correlates with higher ROS production (Galindo-Albarran et al., 2016; Li et al., 2019; Previte et al., 2017). Here, we first considered additional samples and a second ROS sensitive probe to corroborate the higher cROS levels in the neonatal cells (Fig. 1A and B). We then evaluated mROS and identified two cell populations based on its mROS levels (high vs low) (Fig. 1C). We quantified the percentage of cells in the high or low mROS gates from neonatal or adult CD8^+^T cells. We found that neonatal cells have a higher percentage of cells with high mROS, whereas adult cells are more represented in the low mROS gate (Fig. 1D).

**Figure 1.**
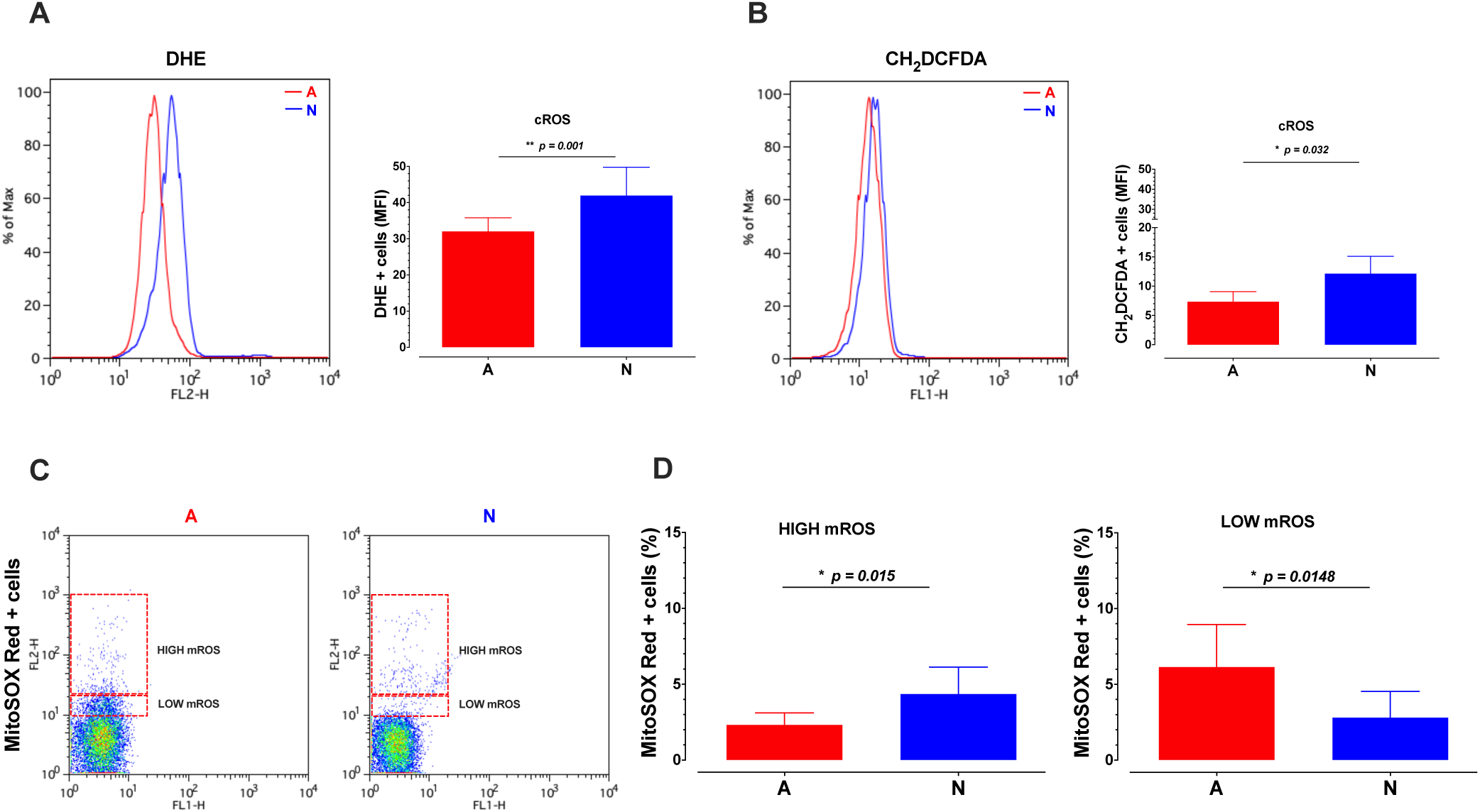
Neonatal CD8^+^T cells have higher basal cROS and mROS levels than their adult counterparts. Purified human CD8^+^T lymphocytes from neonates (N) or adults (A) donors were incubated with 2 μM of DHE (A), 10 μM of Carboxy-H_2_DCFDA (B) or 2.5 μM of the MitoSOX Red (C, and D) fluorescent dyes for 15 minutes. The fluorescence of the cells was assessed using a FACScalibur Flow Cytometer, the graph shows the fluorescence of the cells on the FL-2H channel (DHE, MitoSOX Red) or FL-1H channel (CH_2_DCFDA). At least 4 samples per group were included. An unpaired Mann-Whitney test was used for comparisons between samples from N and A. The level of statistical significance is indicated.

Altogether these results demonstrate that a higher proportion of neonatal CD8^+^T cells display high ROS levels, both in the mitochondria and the cytoplasm.

### T cell activation and mROS

T cell activation induces glycolysis, which could affect the production of ROS. We thus evaluated the changes in ROS after TCR/CD28 cross-linking. Cell activation did not change significantly the cROS levels (Fig. 2A). In the mitochondria, however, activation of adult cells induced a reduction in their low mROS levels, which could be due to the Warburg effect. On the contrary, in the neonatal cells, the stimulation increased the proportion of cells with high mROS (Fig. 2A).

**Figure 2.**
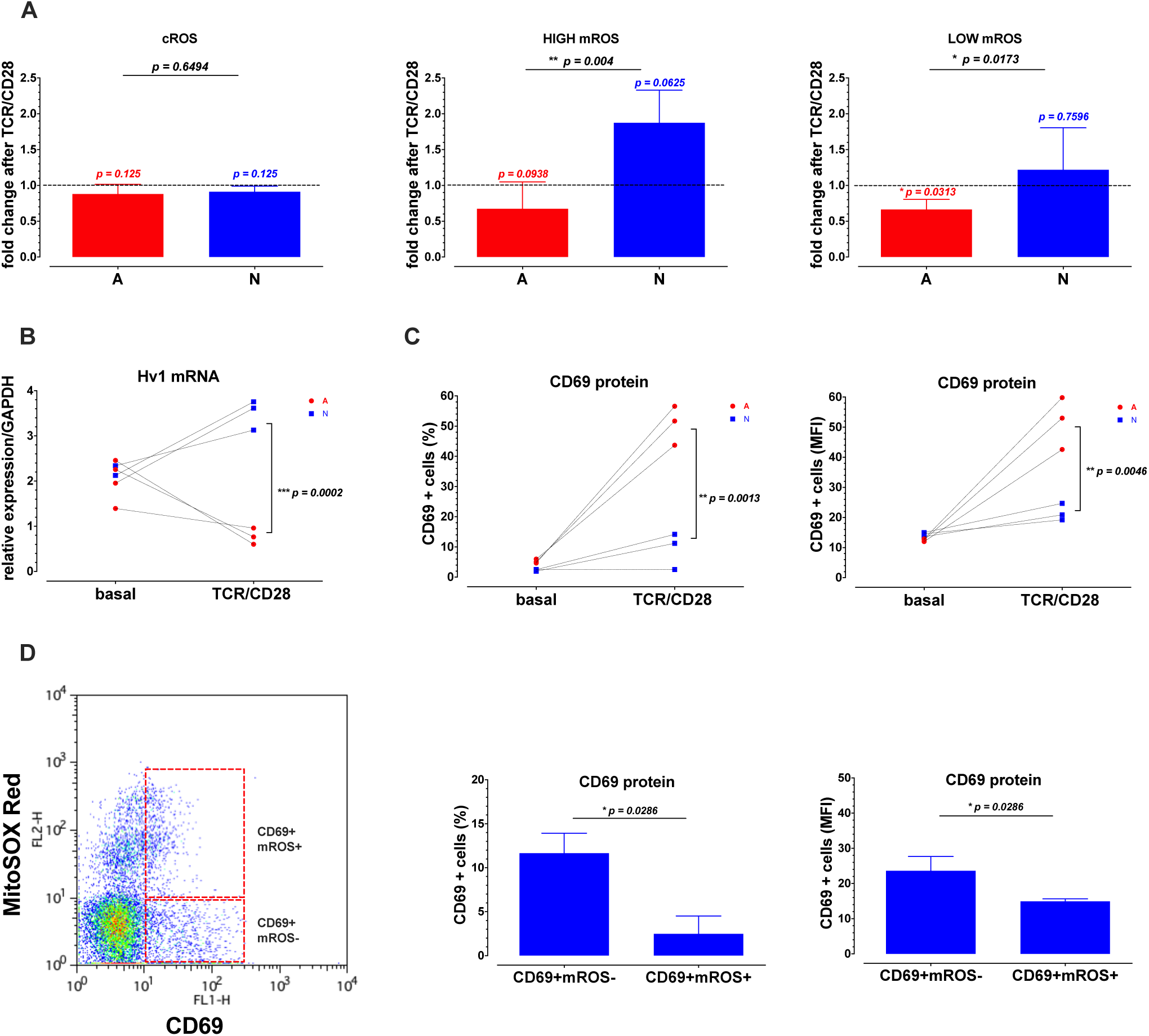
The initial redox stress impairs the activation of neonatal human CD8^+^T cells upon TCR/CD28 stimulation. Purified human CD8^+^T lymphocytes from neonate (N) or adult (A) donors were stimulated with anti-CD3/anti-CD28 antibodies. After 6 hour the cells were collected for measurements. For the ROS measurements cells were incubated with 2 μM of DHE (A) or 2.5 μM of the MitoSOX Red (B and C) fluorescent dyes for 15 minutes. The fluorescence of the cells was assessed using a FACScalibur Flow Cytometer, the graphs A, B and C show the fluorescence or frequency of the cells after normalization to the basal levels. D) Total RNA was extracted for each sample using the TRIzol reagent, the Hv1 proton channel mRNA levels were evaluated using specific primers on a qPCR using the GAPDH mRNA levels as a reference gene. E) The cells were washed and incubated with an anti-CD69 FITC fluorescent antibody for 30 mins the fluorescence or frequency of the cells was analyzed using a FACScalibur Flow Cytometer. F) A MitoSOX Red and anti-CD69 FITC double staining was performed on CD8^+^T cells from neonates. The fluorescence or the frequencies of CD69+mROS- and CD69+mROS+ populations is represented on the bar graphs. At least 4 biological replicates per group were included. An unpaired Mann-Whitney test was used for comparisons between samples from neonates (N) and adult (A) cells, the p-values colored in black denote the significance of these statistical analysis. A Wilcoxon test was used for comparison between paired samples of stimulated N or A groups as compared with the non-stimulated cells, the p-values (red or blue) over the bars denote the significance of these statistical analysis.

The proton channel Hv1 exports protons generated by the NADPH-oxidase enzyme, which controls the excess of reducing power by producing H_2_O_2_, contributing to a high cROS (Asuaje et al., 2017). We measured the expression of Hv1 in neonatal and adult CD8^+^T cells, before and after stimulation. In adult cells, Hv1 expression diminished after activation, while in those of neonates, activation induced Hv1 expression (Fig 2B).

To measure CD8^+^T cell activation, we evaluated CD69 expression. Stimulation induced a higher expression of CD69^+^ cells in adult as compared to neonatal cells. Additionally, in the high mROS gate of the neonatal cells, expression of CD69 was lower as compared to the cells not producing mROS (Fig. 2C).

### Modeling the interplay between T cell activation and mROS

To decipher the interplay between metabolism, ROS and T cell activation, we integrated our own data and literature information on interactions between metabolic pathways and cell signaling in our previous TCR logical model (Rodriguez-Jorge et al., 2019). The resulting model shown in Fig. 3 integrates 111 components and 244 interactions or arcs. This graph includes two inputs corresponding to TCR and CD28 signals. Second messengers, as well as mROS and cROS were also considered, together with key components of the Krebs Cycle, Electron Transport Chain, Fatty acids Oxidation, Glutaminolysis, Pentoses, Fatty Acid Synthesis and Glycolysis pathways. Selected transcription factors link these pathways to output nodes representing cell responses, including Activation, Quiescence (dormant cells), Anergy (incomplete activation of transcription factors) and Metabolic Anergy (unbalanced metabolism, aerobic glycolysis without oxidative mitochondrial metabolism). We also included IL-2 and CD69 as early activation markers. Logical rules specify how each component responds to incoming interactions. The model, including extensive annotations, is provided at the url: http://ginsim.org/node/229 (see also Supp. Table 1).

**Figure 3.**
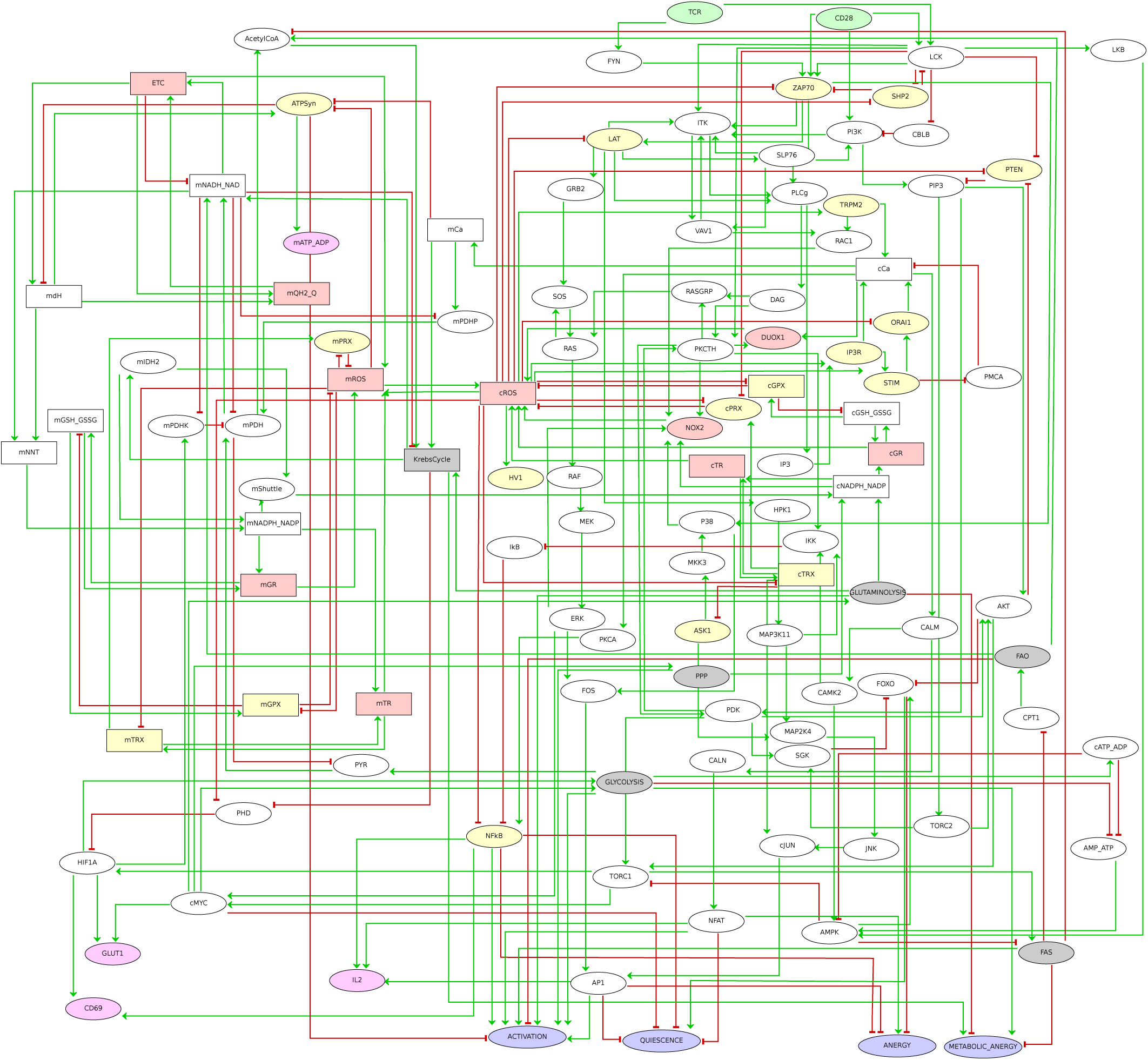
Logical regulatory graph of the TCR-REDOX Metabolism model. The model was designed to study the interaction between the redox state and cellular metabolism and its influence on the activation of human CD8^+^ T lymphocytes. The nodes can be divided into three main categories: mitochondrial nodes (upper left), TCR signaling nodes (upper right), cytosol component nodes (center right and bottom right). A color code further denotes the type of node: green (input), gray (metabolic pathway), pink (ROS or ROS source), yellow (redox sensitive), magenta (output) and purple (phenotype). Elliptical shapes denote Boolean nodes (taking the values 0 or 1), while rectangular shapes denote multilevel (ternary) nodes (taking the values 0, 1 or 2). The arcs connecting the nodes represent positive (green) or negative (red) regulatory influences. In the case of multilevel regulatory nodes, the thresholds required for the activation of the different target nodes can be found in the model available online (http://ginsim.org/node/229).

To evaluate the predictive role of the model, we computed its stable states (Fig. 4A, upper panel). In the absence of TCR or CD28 signal, cells remain quiescent or are kept in metabolic anergy. In the presence of both TCR and CD28 signals, cells could either turn anergic, with high ROS, or undergo activation.

**Figure 4.**
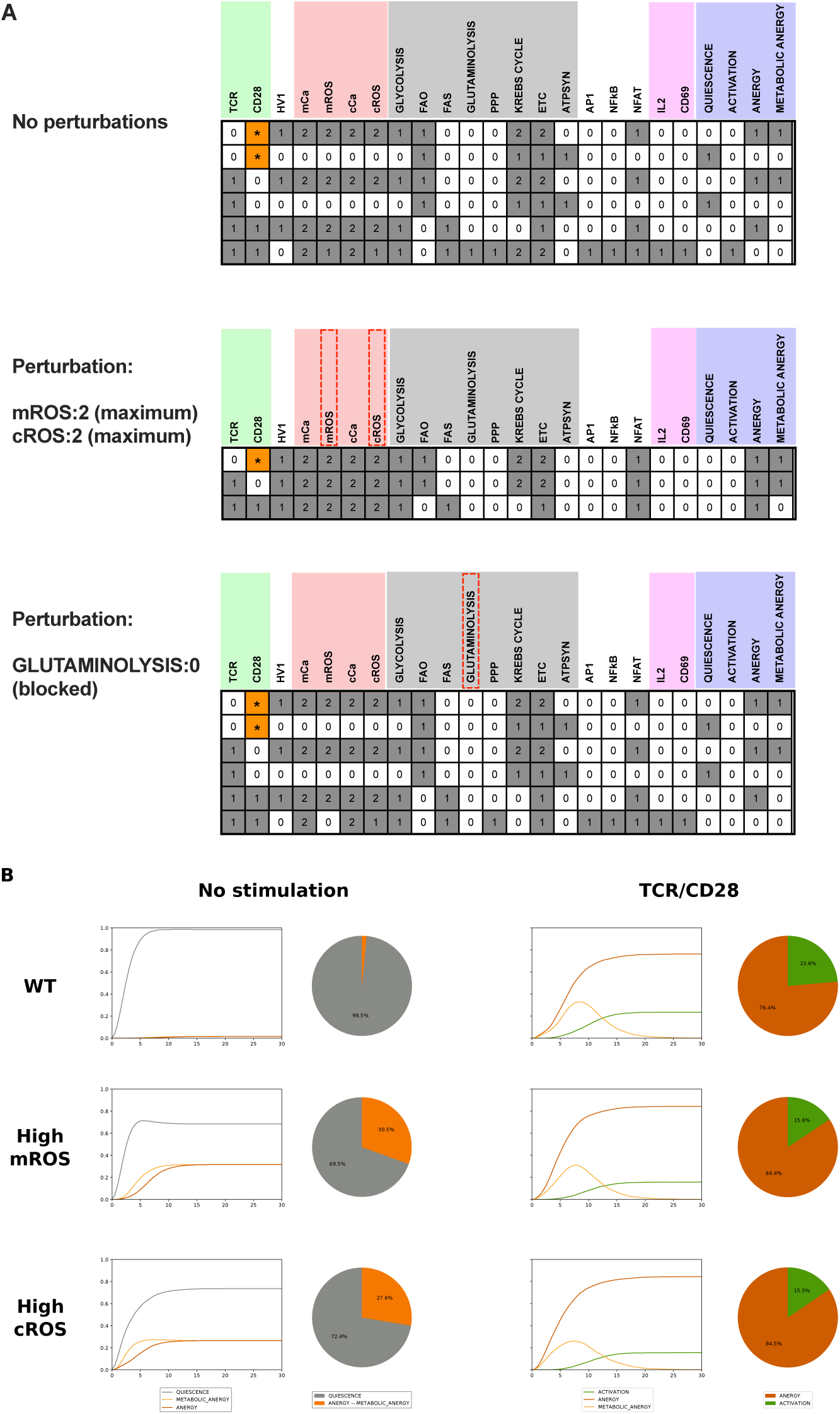
The analysis of the logical model recapitulates the impact of metabolism on T cell activation. (A) Computation of the stable states on selected nodes of the model in three different scenarios (No perturbation; mROS: 2 or cROS: 2 and GLUTAMINOLYSIS: 0). Each row represents a single stable state. For each node, the white cells (0) denote the lowest functional levels, while the gray cells denote intermediate (1) or maximum (2) levels of activity. We further used the MaBoSS software to evaluate the reachability of the different phenotype nodes in the absence of stimulation or upon TCR/CD28 stimulation (B). The pie charts represent the frequency of each phenotype under each condition, while the time plots represent the evolution of each phenotype node over time.

Next, we enforced a fixed high ROS perturbation. In this condition, anergy, metabolic anergy, or a combination of them are predicted (Fig.4A, middle panel).

High arginase levels have been reported in the serum of neonates, which could lead to lower glutamine levels. We thus computed the effect of a blockade of glutaminolysis (Fig.4A, lower panel), which resulted in anergy, metabolic anergy and incomplete activation.

Finally, using the MaBoSS tool, we performed stochastic simulations to assess the reachability of these stable states in unstimulated (Fig.4B) or TCR/CD28 stimulated cells (Fig. 4C). In the non-stimulated (NS) scenario, the majority of the cells (98%) remained quiescent, and only under 2% undergo metabolic anergy. After activation, 23% of cells become active. With fixed activation of either mROS or cROS, 31% or 25% of cells undergo metabolic anergy, respectively. Under these conditions, only 15% of cells reach an active state.

Altogether, our model thus predicts a dramatic effect of ROS status on T cell activation, reproducing our own data, as well as previous observations reviewed in (Simeoni and Bogeski, 2015).

## Discussion

Energy metabolism has a dramatic effect on immune cell homeostasis and response. Neonatal T cells have a unique metabolic and signaling profile, which results in a low response to stimulation.

Neonatal CD8^+^T cells encompass a higher percentage of cells with a very high mROS levels, while low mROS cells are reduced in comparison with adults. Low levels of ROS promote T cell activation, particularly due to the inhibition of phosphatases and the potentiation of NF-κB signaling. High levels of ROS are in contrast detrimental for cell activation (Simeoni and Bogeski, 2015).

The high ROS levels of neonatal cells could be attributed to active glycolysis already in unstimulated cells (Galindo-Albarran et al., 2016) and/or to a low capacity of their mitochondria to control electrons along the Respiratory Chain, producing mROS (Murphy, 2009). Interestingly, cell activation induces Hv1 channel in neonatal but not adult cells, in which its expression is reduced. This channel expels the excess protons generated by the action of the NADPH-oxidase enzyme. Recent evidence suggests that this channel is also located in the internal mitochondrial membrane, contributing to the production and modulation of mROS (Patel et al., 2019). This suggest that neonatal cells cannot modulate the high levels of reducing power produced by glycolysis. A low mitochondrial function of the neonatal cells has been reported (Meszaros et al., 2015). A high level of calcium waves in neonatal cells (Palin et al., 2013) could also convert the mitochondria into calcium storage organelles, which could reduce their metabolic function (Rossi et al., 2019).

Based on our model, we predict that high levels of ROS prevent the activation of PPP, FAS and Glutaminolysis, which are necessary for proper T cell activation. The microenvironment of the neonatal lymphocytes shows high levels of the enzyme arginase (Elahi et al., 2013), which would limit Arginine, which in turn would affect the Glutamine available for de novo synthesis of glutathione, one of the most relevant antioxidant effectors of the cell (Mak et al., 2017). The dynamic analysis of our model indicates that under high ROS levels, over 25% of T cells would be in metabolic anergy, thereby lowering their activation potential, which would tentatively protect newborns from excessive activation at birth, when confronted with many novel antigens.

In conclusion, the metabolic and redox profile of neonatal lymphocytes tentatively impairs their activation potential. This should be addressed in studies focused on boosting neonatal immunity.

## Acknowledgements

We thank Centro Estatal de la Transfusión Sanguínea (Morelos) and Hospital José G. Parres for access to the blood samples. We also thank the mothers of the babies participating on the study, together with all members of the Santana and Thieffry labs. This work was financed by CONACYT Grants 168182 and 257188 and the ECOS/ANUIES/SEP/CONACYT grants M11S01 and M17S02.

## Legends to figures

**Supplementary Table 1. Annotations for the TCR-REDOX-Metabolism Model and specification of the logical rules.** This table has been generated using an export function of the software GINsim and lists the following information for each node of the model (first column):

- a series of database entry identifiers documenting the sources of information used to build the model (second column);

- the Boolean rules defined for each node; note that in the case of multilevel (ternary) nodes, two rules are specified, for values 2, and 1, respectively (third column, upper part of the cells); these rules combine literals (node names) with the standard Boolean operators NOT (denoted by the symbol !), AND (denoted by &), OR (denoted by |), and parentheses whenever required;

- textual annotations explicating the underlying modeling assumptions.

