## Supplemental Table 1 for "An altered metabolism could contribute to the low activation of neonatal CD8^+^T cells"

**Supplementary Table 1. Documentation of the TCR-REDOX-Metabolism model.**

This table has been generated using an export function of the software GINsim and lists the following information for each node of the model (first column):

- a series of database entry identifiers documenting the sources of information used to build the model (second column);
- the Boolean rules defined for each node; note that in the case of multilevel (ternary) nodes, two rules are specified, for values 2, and 1, respectively (third column, upper part of the cells); these rules combine literals (node names) with the standard Boolean operators NOT (denoted by the symbol !), AND (denoted by &), OR (denoted by |), and parentheses whenever required;
- textual annotations explicating the underlying modeling assumptions

| Node ID | References (HGNC, CHEBi or PMID) | Logical rule / **Brief description of the node** |
|---|---|---|
| **TCR** | 1. HGNC:1674<br>2. PMID:19132916 | Input<br>This node represents the TCR heterodimer, including the CD3 complex as well. It is an input node. The functional level 1 is analog to the antigen encounter. The early TCR signaling is reviewed in [2]. |
| **CD28** | 1. HGNC:1653<br>2. PMID:27192564 | Input<br>This node represents the costimulatory CD28 protein. It is an input node. The functional level 1 is analog to the ligation with CD80/CD86 molecule. The relevance of CD28 on the early TCR signaling is reviewed in [2]. |
| **LCK** | 1. HGNC:6524<br>2. PMID:22723799 | (TCR & CD28 & (SHP2 \| !SHP2)) \| (TCR & !CD28 & !SHP2)<br>This node represents the Src-family kinase LCK protein. Its action on the early TCR signaling is reviewed in [2]. |
| **FYN** | 1. HGNC:4037<br>2. PMID:15489916 | TCR<br>This node represents the Src-family kinase FYN protein. Its action on the early TCR signaling is reviewed in [2]. |
| **ZAP70** | 1. HGNC:12858<br>2. PMID:1423621<br>3. PMID:16273097<br>4. PMID:25287889<br>5. PMID:11744042 | (LCK & FYN & CD28 & (!SHP2 \| SHP2) & !cROS) \| (!LCK & FYN & CD28 & !SHP2 & !cROS) \| (LCK & !FYN & CD28 & !SHP2 & !cROS)<br>This node represents the tyrosine kinase ZAP-70, one of the most relevant of the kinases involved on the early TCR signaling [2], [3]. It is a redox sensitive node [4], [5]. |
| **LAT** | 1. HGNC:18874<br>2. PMID:9489702<br>3. PMID:10657671<br>4. PMID:11756537 | ZAP70 & !cROS<br>This node represents the scaffold protein LAT, important for the downstream signaling derived from the TCR activation [2]. It is a redox sensitive node [3], [4]. |
| **SLP76** | 1. HGNC:6529 | LAT & ZAP70 |

| | | |
|---|---|---|
| | 2. PMID:8702662<br>3. PMID:10021361 | This node represents the Src homology 2 (SH2) domain-containing leukocyte protein of 76 kDa, a substrate of ZAP70 connecting TCR signals with downstream activation of RAS and calcium pathways [2], [3]. |
| **ITK** | 1. HGNC:6171<br>2. PMID:11994416<br>3. PMID:9584150<br>4. PMID:15771581 | SLP76 & (VAV1 \| LCK \| ZAP70 \| LAT \| PI3K)<br>This node represents the Tec family kinase ITK. It is an important mediator of the early TCR signals [2], [3], [4]. |
| **VAV1** | 1. HGNC:12657<br>2. PMID:11994416<br>3. PMID:17050525<br>4. PMID:10849438 | ITK & SLP76<br>This node represents the signaling protein VAV1, important in the early TCR signaling events [2], [3], [4]. |
| **RASGRP** | 1. HGNC:9878<br>2. PMID:15899849 | DAG & PKCTH<br>This node represents the RAS exchange factor RASGRP1, an important activator of RAS during TCR activation [2]. |
| **SOS** | 1. HGNC:11187<br>2. PMID:9846483<br>3. PMID:17283063 | GRB2 & RAS<br>This node represents the RAS activator protein SOS. It is an important mediator of the TCR signals [2], [3]. |
| **GRB2** | 1. HGNC:4566<br>2. PMID:9846483<br>3. PMID:8479536 | LAT<br>This node represents the early TCR signaling protein GRB2 involved in the LAT signalosome. |
| **PLCg** | 1. HGNC:9065<br>2. PMID:17148460<br>3. PMID:11994416 | SLP76 & ITK & LAT<br>This node represents the phospholipase C gamma, an important signaling protein connecting the TCR signals with the downstream calcium currents [2], [3]. |
| **PI3K** | 1. HGNC:8975<br>2. PMID:11994416<br>3. PMID:8183372<br>4. PMID:12121659 | (SLP76 & CD28 & (!CBLB \| CBLB)) \| (!SLP76 & CD28 & !CBLB) \| (SLP76 & !CD28 & !CBLB)<br>This node represents the phosphatidylinositol 3'-kinase (PI3K), an important mediator of the CD28 signaling pathway [2], [3]. This node is important for the metabolic shift necessary upon T cell activation [4]. |
| **PIP3** | 1. CHEBI:16618<br>2. PMID:12670391<br>3. PMID:30692200 | PI3K & !PTEN<br>This node represents the second messenger phosphatidylinositol 3,4,5-trisphosphate (PIP3), important in the PI3K signaling pathway [2], [3]. |
| **PDK** | 1. HGNC:8816<br>2. PMID:19122654<br>3. PMID:28152304<br>4. PMID:23183047 | PIP3 \| PKCTH<br>This node represents the enzyme phosphoinositide-dependent kinase 1. It is important for integrating the TCR and CD28 signals [2], [3], [4]. |
| **DAG** | 1. CHEBI:18035<br>2. PMID:17548359 | PLCg<br>This node represents the molecule diacylglycerol, an important second messenger downstream the TCR signaling pathway [2]. |
| **PKCTH** | 1. HGNC:9410<br>2. PMID:17548359<br>3. PMID:17544292 | DAG & LCK & PDK<br>This node represents the kinase PKC theta, a $Ca^{2+}$ independent and DAG dependent PKC [2], [3]. |
| **RAC1** | 1. HGNC:9801<br>2. PMID:15258578<br>3. PMID:30007118<br>4. PMID:24598074 | TRPM2 \| VAV1<br>This node represents the Rho small GTPase RAC. It is an important subunit for the activation of the NADPH-oxidase complex [2], [3], [4]. |

| | | |
|---|---|---|
| **AKT** | 1. HGNC:391<br>2. PMID:12121659<br>3. PMID:18354169<br>4. PMID:15718470<br>5. PMID:28137869<br>6. PMID:29523440<br>7. PMID:29109121 | PIP3 & PDK & TORC2<br>This node represents the serine/threonine kinase Akt. It is an important mediator of the metabolic shift required for T cell activation. |
| **NOX2** | 1. HGNC:2578<br>2. PMID:15258578<br>3. PMID:25081034<br>4. PMID:19028840<br>5. PMID:29757466 | RAC1 & PKCTH & ERK & P38 & cNADPH_NADP<br>This node represents the NADPH-oxidase enzyme complex assembled at the cytoplasmic membrane, which is activated upon TCR activation [2] It is composed of several subunits [4]. It is a source of cROS. |
| **DUOX1** | 1. HGNC:3062<br>2. PMID:20682913 | cCa & PKCTH<br>This node represents the nonphagocytic NADPH oxidase enzime DUOX1. It is a ROS source important for the early TCR signaling [2]. |
| **SHP2** | 1. HGNC:9644<br>2. PMID:17982034<br>3. PMID:20682913 | !(cROS \| LCK)<br>This node represents the tyrosine phosphatase SHP-2, an important negative modulator of the early TCR signaling. It is a redox sensitive node [2], [3]. |
| **PTEN** | 1. HGNC:9588<br>2. PMID:15534200 | !(cROS \| LCK \| AKT)<br>This node represents the phosphatase with sequence homology to tensin (PTEN), a negative modulator of PI3K signaling pathway. It is a redox sensitive node [2]. |
| **CBLB** | 1. HGNC:1542<br>2. PMID:11087752<br>3. PMID:12193687 | !LCK<br>This node represents the E3 ubiquitin ligase Cbl-b, an important negative modulator of the T cell signaling [2], [3]. |
| **RAS** | 1. HGNC:6407<br>2. PMID:10807788<br>3. PMID:9582122<br>4. PMID:15899849<br>5. PMID:9846483<br>6. PMID:17283063 | RASGRP \| SOS<br>This node represents the small GTPase RAS, important for T cells activation [6]. |
| **TRPM2** | 1. HGNC:12339<br>2. PMID:26839633<br>3. PMID:24009611<br>4. PMID:23302782<br>5. PMID:23077651<br>6. PMID:22547068<br>7. PMID:15952035 | cROS<br>This node represents the TRPM2 cation channel located at the cytoplasmic membrane. It is a selective calcium channel [3], sensitive to the redox status of the cell [4], [5], [6]. |
| **PMCA** | 1. HGNC:814<br>2. PMID:22246182<br>3. PMID:14966303 | !STIM<br>This node represents the Plasma Membrane Calcium ATPase. It is a pump that extrudes calcium and modulates its cytoplasmic concentration [2], [3]. |
| **ORAI1** | 1. HGNC:25896<br>2. PMID:20354224<br>3. PMID:19754898<br>4. PMID:16582901 | STIM & !cROS<br>This node represents the ORAI1 calcium channel. It is located at the cytoplasmic membrane and is the main channel responsible for the Store Operated Calcium Entry (SOCE) in T cells [3], [4]. It is a redox sensitive protein [2]. |
| **IP3** | 1. CHEBI 16595 | PLCg |

<table>
<tr><td rowspan="1"></td><td>2. PMID:26052328<br>3. PMID:23805141</td><td>This node represents the second messenger inositol-1,4,5-trisphosphate (IP3), produced by the action of the PLC gamma enzyme. It is an important connection between the TCR signals and the calcium currents [2], [3].</td></tr>
<tr><td rowspan="2">IP3R</td><td rowspan="2">1. HGNC: 6180<br>2. PMID:26052328<br>3. PMID:30228182</td><td>IP3 | cROS</td></tr>
<tr><td>This node represents the inositol trisphosphate receptor (IP3) located at the endoplasmic reticulum membrane. It is a calcium channel opened by the ligation with inositol-1,4,5-trisphosphate (IP3). It induces the depletion of the internal calcium stores [2]. It is a redox sensitive node [3].</td></tr>
<tr><td rowspan="2">STIM</td><td rowspan="2">1. HGNC:11386<br>2. PMID:15866891<br>3. PMID:16005298<br>4. PMID:24053140<br>5. PMID:28900911</td><td>IP3R | cROS</td></tr>
<tr><td>This node represents the calcium sensor and activator of SOCE STIM1, located at the endoplasmic reticulum. It senses the depletion of the internal calcium reserves and activates ORAI1 [2], [3]. It is also a redox sensitive node [4], [5].</td></tr>
<tr><td rowspan="2">CALM</td><td rowspan="2">1. HGNC:1442<br>2. PMID:19290928</td><td>cCa</td></tr>
<tr><td>This node represents calmodulin. It is a sensor for intracellular calcium [2].</td></tr>
<tr><td rowspan="2">CALN</td><td rowspan="2">1. HGNC:9315<br>2. PMID:1715244<br>3. PMID:27941787<br>4. PMID:11015619</td><td>CALM</td></tr>
<tr><td>This node represents the phosphatase calcineurin. This enzyme activates NFAT transcription factor upon elevated cytosolic calcium levels [2], [3], [4].</td></tr>
<tr><td rowspan="2">RAF</td><td rowspan="2">1. HGNC:9829<br>2. PMID:9826444<br>3. PMID:14737111<br>4. PMID:19542438</td><td>RAS</td></tr>
<tr><td>This node represents the serine/threonine kinase Raf-1. It is a MAPKKK upstream of ERK [2], [3], [4].</td></tr>
<tr><td rowspan="2">MEK</td><td rowspan="2">1. HGNC:6847<br>2. PMID:19542438<br>3. PMID:12899942</td><td>RAF</td></tr>
<tr><td>This node represents the mitogen-activated protein kinase kinase 7. It is an activator of ERK [2], [3].</td></tr>
<tr><td rowspan="2">MAP3K11</td><td rowspan="2">1. HGNC:6850<br>2. PMID:10713178<br>3. PMID:10849438</td><td>HPK1</td></tr>
<tr><td>This node represents the Mitogen Activated Kinase Kinase Kinase 11, or MLK3. It is an upstream activator of the NFkB and JNK pathways [2], [3].</td></tr>
<tr><td rowspan="2">MAP2K4</td><td rowspan="2">1. HGNC:6844<br>2. PMID:29248490<br>3. PMID:10785355</td><td>MAP3K11 | ASK1</td></tr>
<tr><td>This node represents the MKK4, a MAPKK important in the activation of JNK in T cells [2], [3].</td></tr>
<tr><td rowspan="2">MKK3</td><td rowspan="2">1. HGNC:6843<br>2. PMID:8974401</td><td>ASK1</td></tr>
<tr><td>This node represents the enzyme MAPKK3. It is a MAPKK involved in the activation of p38 during intracellular stress [2].</td></tr>
<tr><td rowspan="2">ASK1</td><td rowspan="2">1. HGNC:6857<br>2. PMID:18206122<br>3. PMID:11274345<br>4. PMID:17883330</td><td>!cTRX</td></tr>
<tr><td>This node represents the Apoptosis signal-regulating kinase 1 (ASK1). It is a MAPKKK upstream of JNK and P38 which activates in response to cellular stress. It is a redox sensitive node [2], [3], [4].</td></tr>
<tr><td rowspan="2">HPK1</td><td rowspan="2">1. HGNC:6863<br>2. PMID:10849438</td><td>LAT</td></tr>
<tr><td>This node represents the hematopoietic progenitor kinase 1 (HPK1), a serine/threonine kinase important for the signaling downstream the TCR</td></tr>
</table>

| | | |
|---|---|---|
| | | [2]. |
| **HV1** | 1. HGNC:28240<br>2. PMID:28013412<br>3. PMID:23798303<br>4. PMID:24890927<br>5. PMID:20139987<br>6. PMID:25425665<br>7. PMID:29643227<br>8. PMID:31060879 | cROS<br><br>This node represents the Hv1 proton channel located at the cytoplasmic membrane [3]. This channel extrudes the excess of protons generated by the NADPH-oxidase enzyme, thus preventing cellular acidification [2]. Recent evidences support a role in the modulation of the mROS generation located at the internal mitochondrial membrane [8]. |
| **mCa (multilevel)** | 1. CHEBI:29108<br>2. PMID:20668216<br>3. PMID:30622345<br>4. PMID:29032101<br>5. PMID:30982525<br>6. PMID:26100311<br>7. PMID:29030115 | 2 -cCa:2<br>1- cCa:1<br><br>This node represents the mitochondrial level of the Ca ion, an important second messenger for T cells signaling [3]. It is important for the mitochondrial metabolic activity [4], [5], [7]. Recent evidences suggest that the Ca levels are differentially regulated between T cells from neonates and adults [6]. It is a multilevel node, taking<br>level 2 iff very high $mCa^{2+}$ levels;<br>level 1iff intermediate $mCa^{2+}$ levels;<br>level 0 iff very low $mCa^{2+}$ levels. |
| **mROS (multilevel)** | 1. CHEBI:26523<br>2. PMID:23415911<br>3. PMID:23880762<br>4. PMID:30466061<br>5. PMID:30057684<br>6. PMID:19061483<br>7. PMID:27965578<br>8. PMID:26100311 | 2 - (!(mPRX \| mGPX) & (cROS \| ETC \| mGR \| mTR)) \| (cROS & (mPRX \| mGPX \| ETC \| mGR \| mTR))<br>1 - ((!mPRX & mGPX) \| (mPRX & !mGPX)) & (ETC \| mGR \| mTR) & !cROS<br><br>This node represents the levels of Reactive Oxigen Species generated in the mitochondria (mROS). This is representing mainly by hydrogen peroxide (H2O2), but it also includes the superoxide anion (O2.-) and other derivatives. It is an important second messenger for signaling but it is detrimental at higher levels. Thus, its concentration is maintained under a strict balance between production and scavenging. It is a multilevel node taking<br>level 2 iff very high damaging levels of mROS;<br>level 1 iff normal signaling levels of mROS;<br>level 0 iff very low levels of mROS. |
| **mQH2_Q (multilevel)** | 1. CHEBI:17976<br>2. PMID:19061483<br>3. PMID:27085844 | 2 - ETC & mdH<br>1 - ETC & !mdH<br><br>This node represents the ratio between the reduced ($QH_2$) and the oxidized (Q) ubiquinone, the electron carrier in the mitochondrial internal membrane. At some conditions of very reduced values, it is a source of mROS [2], [3]. It is a multilevel node taking<br>level 2 iff very reduced;<br>level 1 iff normal physiological levels;<br>level 0 iff very oxidized. |
| **mNADH_NAD (multilevel)** | 1. CHEBI:16908<br>2. PMID:28648096 | 2 - (KrebsCycle \| mPDH \| FAO) & !ETC<br>1 - (KrebsCycle \| mPDH \| FAO) & ETC |

| | | |
|---|---|---|
| | 3. PMID:20175987<br>4. PMID:23442855<br>5. PMID:19061483 | This node represents the ratio between the redox couple of the reduced (NADH) vs oxidized ($NAD^+$) coenzyme at the mitochondria. It is an important indicator of the energetic status of the cell [2], [3], [4], [5]. It is a multilevel node taking<br>level 2 iff corresponds to a very high NADH/$NAD^+$ ratio, the couple is at an extreme of reduction;<br>level 1 iff corresponds to a high or medium NADH/$NAD^+$ ratio, the couple is at normal physiological levels of reduction;<br>level 0 iff corresponds to a very low NADH/$NAD^+$ ratio, the couple is at an extreme of oxidation. |
| **mGSH_GSSG (multilevel)** | 1. CHEBI:16856<br>2. PMID:25701705<br>3. PMID:28558477 | 2 - mGR & !mGPX<br>1 - mGR & mGPX |
| | | This node represents the ratio between the reduced (GSH) and the oxidized (GSSG) mitochondrial pool of glutathione, an important antioxidant of the cell [3]. In some cases of reductive stress, the extremely reduced GSH/GSSG ratio is an inductor of ROS production by the enzyme Glutathione Reductase [2]. It is a multilevel node taking<br>level 2 iff highly reduced;<br>level 1 iff normal physiological reduction status;<br>level 0 iff highly oxidized. |
| **mdH (multilevel)** | 1. CHEBI:24636<br>2. PMID:29626541<br>3. PMID:19061483 | 2 - ETC & !ATPSyn<br>1 - ETC & ATPSyn |
| | | This node represents the proton concentration at the mitochondrial intermediate membrane space. It is the proton motive force which drive ATP production by the ATP synthase. Several evidences connect the proton motive force to the generation on mROS [2], [3]. It is a multilevel node taking<br>level 2 iff very high proton motive force;<br>level 1 iff intermediate levels of proton motive force;<br>level 0 iff very low proton motive force. |
| **mNADPH_NADP (multilevel)** | 1. CHEBI:16474<br>2. PMID:28648096<br>3. PMID:20175987<br>4. PMID:23442855 | 2 - (mNNT:2 & !mIDH2) \| (mNNT:2 & mIDH2) \| (mNNT:1 & mIDH2)<br>1 - (!mNNT & mIDH2) \| (mNNT:1 & !mIDH2) |
| | | This node represents the ratio between the redox couple of the reduced (NADHP) vs oxidized ($NADP^+$) coenzyme in the mitochondria. It is an important indicator of the reducing power of the cell [2], [3], [4]. It is a multilevel node taking<br>level 2 for a very high NADPH/$NADP^+$ ratio, the couple is at an extreme of reduction<br>level 1 for a high or medium NADPH/$NADP^+$ ratio, the couple is at normal physiological levels of reduction.<br>level 0 for a very low NADPH/$NADP^+$ ratio, the couple is at an extreme of oxidation. |
| **mNNT (multilevel)** | 1. HGNC:7863<br>2. PMID:28648096<br>3. PMID:16730324 | 2 - (mNADH_NAD:2 & mdH:2) \| (mNADH_NAD:1 & mdH:2) \| (mNADH_NAD:2 & mdH:1)<br>1 - mNADH_NAD:1 & mdH:1 |

| | | |
|---|---|---|
| | 4. PMID:20007326 | This node represents the NAM nucleotide transhydrogenase, an important mediator of the conversion between NADH and NADPH pools in the mitochondria. It is regulated by the proton motive force. It is a multilevel node. |
| **mIDH2** | 1. HGNC:5383<br>2. PMID:28648096<br>3. PMID:20007326 | KrebsCycle |
| | | This node represents the mitochondrial enzyme Isocitrate dehydrogenase II. It is a source of mitochondrial NADPH [2], [3]. |
| **mGPX (multilevel)** | 1. HGNC:4553<br>2. PMID:23397885 | 2 - mGSH_GSSG:2 & !mROS<br>1 - mGSH_GSSG:1 & !mROS |
| | | This node represents the mitochondrial levels of the glutathione peroxidase. It is an important element of the antioxidant machinery [2]. It is a multilevel node. |
| **mGR (multilevel)** | 1. HGNC:4623<br>2. PMID:25701705 | 2 - mGSH_GSSG:2 & (mNADPH_NADP:1 \| mNADPH_NADP:2)<br>1 - (mNADPH_NADP:1 & !mGSH_GSSG) \| (mNADPH_NADP:1 & mGSH_GSSG:1) \| (mNADPH_NADP:2 & !mGSH_GSSG) \| (mNADPH_NADP:2 & mGSH_GSSG:1) |
| | | This node represents the mitochondrial enzyme Glutathione Reductase (mGR). This enzyme is important in the generation of reduced Glutathione from NADPH. It is a source of ROS at extreme reduction conditions. It is a multilevel node [2] taking<br>level 2 iff highly reduced enzyme;<br>level 1 iff normal physiological redox status of the enzyme;<br>level 0 iff highly oxidized enzyme. |
| **mPDHP** | 1. HGNC:9279<br>2. PMID:30982525 | mCa & !mNADH_NAD |
| | | This node represents the enzyme Pyruvate Dehydrogenase Phosphatase located at the mitochondria. This enzyme is a positive regulator of the Pyruvate Dehydrogenase. It is an important mediator of the oxidative metabolism in the mitochondria [2]. |
| **mTR (multilevel)** | 1. HGNC:18155<br>2. PMID:29278740<br>3. PMID:25701705<br>4. PMID:20175987 | 2 - mTRX:2 & mNADPH_NADP:2<br>1 - (mNADPH_NADP:1 & !mTRX) \| (mNADPH_NADP:1 & mTRX:1) \| (mNADPH_NADP:2 & !mTRX) \| (mNADPH_NADP:2 & mTRX:1) \| (mNADPH_NADP:1 & mTRX:2) |
| | | This node represents the mitochondrial enzyme Thioredoxin Reductase (mTR) [2]. This enzyme is important in the generation of reduced Thioredoxin from NADPH. It is a source of ROS at extreme reduction conditions [3], [4]. It is a multilevel node taking<br>level 2 iff highly reduced enzyme;<br>Level 1 iff normal physiological redox status of the enzyme;<br>Level 0 iff highly oxidized enzyme. |
| **mTRX (multilevel)** | 1. HGNC:17772<br>2. PMID:23397885 | 2 - mTR & !mROS<br>1 - mTR & mROS |

| | | |
|---|---|---|
| | 3. PMID:25701705 | This node represents the thioredoxin mitochondrial pool. This is an important element on the anti-oxidant machinery of the cell. It receives electrons from NADPH and transfers the electrons to other signaling proteins [2]. In some cases of reductive stress, the extremely reduced TRX pool is an inductor of ROS production by the enzyme Thioredoxin Reductase [3]. It is a multilevel node taking<br>level 2 iff highly reduced;<br>level 1 iff normal physiological reduction status;<br>level 0 iff highly oxidized. |
| **mPRX** | 1. HGNC:9354<br>2. PMID:23397885 | (mTRX & !mROS) \| (mTRX:2 & mROS)<br><br>This node represents the mitochondrial pool of peroxiredoxin. It is an important antioxidant protein also involved in redox signaling [2]. |
| **mShuttle** | 1. HGNC:10979<br>2. PMID:28648096<br>3. PMID:20007326 | mIDH2 & mNADPH_NADP<br><br>This node represents the 2-oxoglutarate/isocitrate NADP(H)-redox shuttle located in the mitochondria. It connects the mitochondrial and cytosolic NADPH pools [2], [3]. |
| **mPDH** | 1. HGNC:8806<br>2. PMID:30982525 | (mPDHP \| PYR) & !mPDHK & !mNADH_NAD<br><br>This node represents the Pyruvate Dehydrogenase enzyme, located in the mitochondria. This enzyme is important for the complete oxidation of glucose derivatives [2]. |
| **mPDHK** | 1. HGNC:8809<br>2. PMID:16517405<br>3. PMID:25437876 | !mNADH_NAD & HIF1A<br><br>This node represents the enzyme Pyruvate Dehydrogenase Kinase located at the mitochondria. This enzyme is a negative regulator of the Pyruvate Dehydrogenase. It is an important mediator of the metabolic shift towards aerobic glycolysis [2], [3]. |
| **mATP_ADP** | 1. CHEBI:15422<br>2. PMID:25229666 | ATPSyn<br><br>This node represents the mitochondrial ATP/ADP ratio. It is an indicator of the oxidative metabolism of quiescent cells supported mainly by the mitochondria, different from the Warburg metabolism of proliferative cells supported mainly by the cytosol [2]. |
| **cCa (multilevel)** | 1. CHEBI:29108<br>2. PMID:29030115<br>3. PMID:30622345<br>4. PMID:26100311<br>5. PMID:23408835 | 2 - (IP3R & ORAI1 & TRPM2 & !PMCA) \| (IP3R & ORAI1 & !TRPM2 & !PMCA) \| (IP3R & !ORAI1 & TRPM2 & !PMCA)<br>1 - (IP3R & !ORAI1 & !TRPM2 & !PMCA) \| (!IP3R & !ORAI1 & TRPM2 & !PMCA) \| (!IP3R & ORAI1 & TRPM2 & !PMCA) \| (!IP3R & ORAI1 & !TRPM2 & !PMCA)<br><br>This node represents the cytosolic levels of the $Ca^{2+}$ ion, an important second messenger for T cells signaling [2], [3]. Recent evidences suggest that the $Ca^{2+}$ levels are differentially regulated between T cells from neonates and adults [4], [5]. It is a multilevel node taking<br>level 2 iff very high $mCa^{2+}$ levels;<br>level 1 iff intermediate $mCa^{2+}$ levels;<br>level 0 iff very low $mCa^{2+}$ levels. |

| | | |
|---|---|---|
| **cROS (multilevel)** | 1. CHEBI:26523<br>2. PMID:17982034<br>3. PMID:27965578 | 2 - (!(cPRX \| cGPX) & (NOX2 \| mROS \| DUOX1 \| cTR \| cGR)) \| (mROS & (cPRX \| cGPX \| NOX2 \| DUOX1 \| cTR \| cGR))<br>1 - ((cPRX & !cGPX) \| (!cPRX & cGPX)) & (NOX2 \| DUOX1 \| cTR \| cGR) & !mROS |
| | | This node represents the levels of Reactive Oxigen Species generated in the cytosol (cROS). This is representing mainly by hydrogen peroxide ($H_2O_2$), but it also includes the superoxide anion ($O_2^{\cdot -}$) and other derivatives. It is an important second messenger for signaling but it is detrimental at higher levels. Thus, its concentration is maintained under a strict balance between production and scavenging. It is a multilevel node taking<br>level 2 iff very high damaging levels of cROS;<br>level 1 iff normal signaling levels of cROS;<br>level 0 iff very low levels of cROS. |
| **cNADPH_NADP (multilevel)** | 1. CHEBI:16474<br>2. PMID:28648096<br>3. PMID:20175987<br>4. PMID:23442855 | 2 - (mShuttle & PPP & GLUTAMINOLYSIS) \| (!mShuttle & PPP & GLUTAMINOLYSIS) \| (mShuttle & !PPP & GLUTAMINOLYSIS) \| (mShuttle & PPP & !GLUTAMINOLYSIS)<br>1 - (mShuttle & !PPP & !GLUTAMINOLYSIS) \| (!mShuttle & PPP & !GLUTAMINOLYSIS) \| (!mShuttle & !PPP & GLUTAMINOLYSIS) |
| | | This node represents the ratio between the redox couple of the reduced (NADHP) vs oxidized ($NADP^+$) coenzyme at the cytosol. It is an important indicator of the reducing power of the cell [2], [3], [4]. It is a multilevel node taking<br>level 2 iff corresponds to a very high $NADPH/NADP^+$ ratio, the couple is at an extreme of reduction;<br>level 1 iff corresponds to a high or medium $NADPH/NADP^+$ ratio, the couple is at normal physiological levels of reduction;<br>level 0 iff corresponds to a very low $NADPH/NADP^+$ ratio, the couple is at an extreme of oxidation. |
| **cGSH_GSSG (multilevel)** | 1. CHEBI:16856<br>2. PMID:30232291<br>3. PMID:25701705 | 2 - cGR & !cGPX<br>1 - cGR & cGPX |
| | | This node represents the ratio between the reduced (GSH) and the oxidized (GSSG) cytosolic pool of glutathione, an important antioxidant of the cell [2]. In some cases of reductive stress, the extremely reduced GSH/GSSG ratio is an inductor of ROS production by the enzyme Glutathione Reductase [3]. It is a multilevel node taking<br>level 2 iff highly reduced;<br>level 1 iff normal physiological reduction status;<br>level 0 iff highly oxidized. |
| **cATP_ADP** | 1. CHEBI:15422<br>2. PMID:20814441 | GLYCOLYSIS |
| | | This node represents the cytosol ATP/ADP ratio. It is an indicator of the Warburg metabolism of proliferative cells supported mainly by the cytosol, different from the oxidative metabolism of quiescent cells supported mainly by the mitochondria [2]. |
| **AMP_ATP** | 1. CHEBI:16027 | !cATP_ADP & !GLYCOLYSIS |

| | | |
|---|---|---|
| | 2. PMID:20814441 | This node represents the cellular AMP to ATP ratio. An important indicator of the energy status of the cell [2]. |
| **cTRX (multilevel)** | 1. HGNC:12435<br>2. PMID:29749372<br>3. PMID:23397885 | 2 - cTR & !cROS<br>1 - cTR & cROS:1 |
| | | This node represents the thioredoxin cytosolic pool. This is an important element on the antioxidant machinery of the cell. It receives electrons from NADPH and transfers the electrons to other signaling proteins [2]. In some cases of reductive stress, the extremely reduced TRX pool is an inductor of ROS production by the enzyme Thioredoxin Reductase [3]. It is a multilevel node taking<br>level 2 iff highly reduced;<br>level 1 iff normal physiological reduction status;<br>level 0 iff highly oxidized. |
| **cGR (multilevel)** | 1. HGNC:4623<br>2. PMID:25701705 | 2 - cGSH_GSSG:2 & (cNADPH_NADP:1 \| cNADPH_NADP:2)<br>1 - (cNADPH_NADP:1 & !cGSH_GSSG) \| (cNADPH_NADP:1 & cGSH_GSSG:1) \| (cNADPH_NADP:2 & !cGSH_GSSG) \| (cNADPH_NADP:2 & cGSH_GSSG:1) |
| | | This node represents the cytoplasmic enzyme Glutathione Reductase (cGR). This enzyme is important in the generation of reduced Glutathione from NADPH. It is a source of ROS at extreme reduction conditions. It is a multilevel node [2] taking<br>level 2 iff highly reduced enzyme;<br>level 1 iff normal physiological redox status of the enzyme;<br>level 0 iff highly oxidized enzyme. |
| **cGPX (multilevel)** | 1. HGNC:4554<br>2. PMID:23397885 | 2 iff cGSH_GSSG:2 & !cROS<br>1 iff cGSH_GSSG:1 & !cROS |
| | | This node represents the cytoplasmic levels of the glutathione peroxidase. It is an important element of the antioxidant machinery [2]. It is a multilevel node. |
| **cTR (multilevel)** | 1. HGNC:12437<br>2. PMID:29278740<br>3. PMID:25701705<br>4. PMID:20175987 | 2 - cTRX:2 & cNADPH_NADP:2<br>1 - (cNADPH_NADP:1 & !cTRX) \| (cNADPH_NADP:1 & cTRX:1) \| (cNADPH_NADP:2 & !cTRX) \| (cNADPH_NADP:2 & cTRX:1) \| (cNADPH_NADP:1 & cTRX:2) |
| | | This node represents the cytosolic enzyme Thioredoxin Reductase (cTR) [3]. This enzyme is important in the generation of reduced Thioredoxin from NADPH. It is a source of ROS at extreme reduction conditions [4], [2]. It is a multilevel node taking<br>level 2 iff highly reduced enzyme;<br>level 1 iff normal physiological redox status of the enzyme;<br>level 0 iff highly oxidized enzyme. |
| **cPRX** | 1. HGNC:9354<br>2. PMID:23397885 | cTRX & !(cROS & LCK) |
| | | This node represents the cytosolic pool of peroxiredoxin. It is an important antioxidant protein also involved in redox signaling [2]. |

| | | |
|---|---|---|
| **TORC1** | 1. HGNC:3942<br>2. PMID:23183047<br>3. PMID:24076634<br>4. PMID:21233853<br>5. PMID:14668532<br>6. PMID:30320109<br>7. PMID:25893604<br>8. PMID:30619240<br>9. PMID:22517423 | (PDK \| AKT) & !AMPK<br><br>This node represents the mTORC1 complex. It is an important regulator of the metabolic adaptation of T cells upon activation. |
| **TORC2** | 1. HGNC:3942<br>2. PMID:21310961<br>3. PMID:30320109<br>4. PMID:25893604<br>5. PMID:22517423 | PIP3<br><br>This node represents the mTORC2 complex. It is an important regulator of the metabolic adaptation of T cells upon activation. |
| **SGK** | 1. HGNC:10810<br>2. PMID:11154281<br>3. PMID:22517423 | PDK & TORC2<br><br>This node represents the Serum and glucocorticoid inducible kinase. It is activated by mTORC2 complex and is important in the inhibition of the FOXO1 transcription factor during T cell activation [2], [3]. |
| **LKB** | 1. HGNC:11389<br>2. PMID:21930968<br>3. PMID:21487392 | LCK<br><br>This node represents the serine/threonine kinase liver kinase B 1 (LKB1). It is an important regulator of T cell activation and metabolism [2], [3]. |
| **AMPK** | 1. HGNC:9376<br>2. PMID:21670147<br>3. PMID:18719600<br>4. PMID:16818670<br>5. PMID:23310952<br>6. PMID:21892142<br>7. PMID:25360847 | (LKB \| CAMK2 \| AMP_ATP) & !cATP_ADP<br><br>This node represents the AMP-activated protein kinase (AMPK). This is an important regulator of cellular energetic metabolism. It controls the transition between the basal metabolism of quiescent cells and the highly anabolic metabolism of proliferating cells. |
| **CAMK2** | 1. HGNC:1461<br>2. PMID:15843557 | CALM<br><br>This node represents the calcium/calmodulin-dependent protein kinase II. This enzyme connects the calcium waves with other signaling events important for T cells activation and proliferation [2]. |
| **PHD** | 1. HGNC:8548<br>2. PMID:20199358<br>3. PMID:24491179<br>4. PMID:23535595 | !cROS & !KrebsCycle<br><br>This node represents the enzyme prolyl 4-hydroxylase. It inhibits HIF1 activation in response to high ROS levels or high succinate levels [2], [3], [4]. |
| **IKK** | 1. HGNC:5960<br>2. PMID:10713178<br>3. PMID:10733597 | PKCTH \| CAMK2 \| MAP3K11<br><br>This node represents the I kappa B kinase (IKK). This kinase is activated by several other kinases (PKC theta, CAMK2 or MAP3K11) and inhibits by phosphorylation the IkB inhibitor of the NFkB transcription factor [2], [3]. |
| **IkB** | 1. HGNC:7797<br>2. PMID:7594468 | !IKK<br><br>This node represents I kappa B alpha, a negative modulator of NFkB transcription factor [2]. |
| **PKCA** | 1. HGNC:9393<br>2. PMID:26826124<br>3. PMID:1388136<br>4. PMID:30048879 | cCa<br><br>This node represents the Ca (2+)-dependent protein kinase C A (PKCA). It activates NFkB transcription factor [2], [3], [4]. |
| **CPT1** | 1. HGNC:2328 | !FAS |

| | | |
|---|---|---|
| | 2. PMID:30619240<br>3. PMID:22206904<br>4. PMID:25001241 | This node represents the enzyme carnitine palmitoyl transferase, an important regulatory enzyme controlling the rate of Fatty Acid Oxidation (FAO) in the mitochondria [2], [3], [4]. |
| **PYR** | 1. CHEBI:15361<br>2. PMID:30619240 | GLYCOLYSIS & !mPDH<br>This node represents pyruvate. This is a metabolite derived from the catabolism of glucose through glycolysis [2]. |
| **AcetylCoA** | 1. CHEBI:15351<br>2. PMID:30619240<br>3. PMID:25703630 | (mPDH \| FAO) & !FAS<br>This node represents the mitochondrial pool of the Acetyl CoA metabolite. It is an intermediate metabolite between lipids and carbohydrates. It is a common indicator of the energy fitness of the cell [2], [3]. |
| **GLUT1** | 1. HGNC:11005<br>2. PMID:30619240 | HIF1A \| cMYC<br>This node represents the glucose transporter Glut1, relevant for the glucose uptake required after TCR activation [2]. |
| **GLYCOLYSIS** | 1. 12121659<br>2. PMID:30619240 | HIF1A \| cMYC<br>This node represents the pathway Glycolysis, with enzymes located at the cytosol, responsible for the glucose catabolism. This pathway is upregulated upon TCR activation and is a metabolic signature of highly proliferative cells [1], [2]. |
| **FAO** | 1. PMID:19494812 | CPT1<br>This node represents the Fatty Acid Oxidation (FAO) metabolic pathway rate. This pathway is maintained high during T cell quiescence and is downregulated upon TCR activation [1]. |
| **FAS** | 1. PMID:24567531<br>2. PMID:25592731 | TORC1 & !AMPK<br>This node represents the Fatty Acid Synthesis pathway, regulated mainly by the ACC enzyme [1]. The upregulation of this pathway is important to the cell growth and proliferation after the TCR activation. |
| **GLUTAMINOLYSIS** | 1. PMID:22195744<br>2. PMID:20554958 | cMYC<br>This node represents the rate of the Glutaminolysis metabolic pathway. It is a catabolic pathway upregulated during T cell activation and proliferation [1], [2]. |
| **PPP** | 1. PMID:22195744 | cMYC<br>This node represents the Pentose Phosphate Pathway, important in the generation of reduction power and carbon skeletons for anabolism and T cell growth and proliferation [1]. |
| **KrebsCycle (multilevel)** | 1. PMID:30982525<br>2. PMID:22195744 | KrebsCycle:2 ☐ mCa & (AcetylCoA \| GLUTAMINOLYSIS) & !mNADH_NAD<br>KrebsCycle:1 ☐ (AcetylCoA \| GLUTAMINOLYSIS) & !mCa & !mNADH_NAD<br>This node represents the enzymes of the TCA cycle located at the mitochondrial matrix. It is an important indicator of the energetic status of the cell. It is a multilevel node. |
| **ETC** | 1. PMID:16818739<br>2. PMID:22206904<br>3. PMID:12471050 | 2 - mNADH_NAD & mQH2_Q<br>1 - (!mNADH_NAD & mQH2_Q) \| (mNADH_NAD:1 & !mQH2_Q) \| (mNADH_NAD:2 |

| | | |
|---|---|---|
| | 4. PMID:19061483<br>5. PMID:27085844<br>6. PMID:30982525 | & !mQH2_Q)<br><br>This node represents the enzymes of the Electron Transport Chain located in the internal mitochondrial membrane, which transfer electrons from reduced coenzymes to $O_2$. Several evidences point to the importance of this pathway on T cell activation [2], [3]. At some metabolic situations it generates ROS [4], [5].<br>It is a multilevel node. The ROS production occurs at level 2. |
| **ATPSyn** | 1. HGNC: 823<br>2. PMID:22889218<br>3. PMID:28522037<br>4. PMID:23312134<br>5. PMID:19061483<br>6. PMID:27085844 | mdH & !mROS & !mCa<br><br>This node represents the ATP synthase enzyme complex, located at the internal mitochondrial membrane. It is a redox sensitive node [4], [5], [6]. |
| **P38** | 1. HGNC:6876<br>2. PMID:12670401<br>3. PMID:16542479<br>4. PMID:25151490<br>5. PMID:18451303<br>6. PMID:10924852<br>7. PMID:16799472<br>8. PMID:15735648 | MKK3 \| ZAP70<br><br>This node represents P38, a MAPK important in the activation of T cells. |
| **ERK** | 1. HGNC:6871<br>2. PMID:19017950<br>3. PMID:26990855 | MEK<br><br>This node represents the Extracellular regulated Kinase, a MAPK. It is an important regulator of T cells activation and proliferation [2], [3]. |
| **FOS** | 1. HGNC:3796<br>2. PMID:12972619<br>3. PMID:15708845 | ERK \| P38<br><br>This node represents the subunit FOS, a component of the AP1 transcription factor. |
| **cJUN** | 1. HGNC:6204<br>2. PMID:25680272<br>3. PMID:10970869 | JNK & cTRX<br><br>This node represents the subunit JUN, a component of the AP1 transcription factor. |
| **JNK** | 1. HGNC:6881<br>2. PMID:24673683<br>3. PMID:12670401<br>4. PMID:10924852 | MAP2K4<br><br>This node represents the c-Jun N-terminal kinase. It is a MAPK important in the activation of the AP1 transcription factor. |
| **HIF1A** | 1. HGNC:4910<br>2. PMID:23183047<br>3. PMID:24076634 | TORC1 \| !PHD<br><br>This node represents the hypoxia-inducible factor 1 alpha protein. It is an important mediator of the metabolic shift towards aerobic glycolysis in T cells during activation [2], [3]. |
| **cMYC** | 1. HGNC:7553<br>2. PMID:21704229<br>3. PMID:22195744<br>4. PMID:30619240<br>5. PMID:8035827 | TORC1 & ERK<br><br>This node represents the transcription factor MYC. It is an important regulator of the metabolic shift required for T cells activation and proliferation. |
| **FOXO** | 1. HGNC:3821<br>2. PMID:18391970<br>3. PMID:19658095 | !AKT & !SGK & AMPK<br><br>This node represents the Forkhead transcription factor 1. This factor is important in the homeostatic survival of naive T cells [2], [3]. |
| **AP1** | 1. HGNC:6204 | FOS & cJUN |

| | | |
|---|---|---|
| | 2. HGNC:3796<br>3. PMID:25680272<br>4. PMID:10970869 | This node represents the transcription factor AP1. It is composed of the subunits FOS and JUN. It is an important transcriptional regulator of T cell fate upon stimulation. In combination with AP1 it promotes activation, but alone it promotes anergy [3], [4]. |
| **NFKB** | 1. HGNC:9955<br>2. PMID:26826124<br>3. PMID:21199863<br>4. PMID:11714266 | !(cROS \| IkB) & PKCA<br>This node represents the transcription factor NFkB. It is a combination of multiple heterodimers of p50, p65 and c-Rel. It is an important factor for the T cell activation upon stimulation or antigen encounter [2], [3], [4]. |
| **NFAT** | 1. HGNC:7775<br>2. PMID:20725108<br>3. PMID:12975316<br>4. PMID:25680272 | CALN<br>This node represents the Nuclear Factor of Activated T cells. It is an important transcriptional regulator of T cell fate upon stimulation [2], [3]. In combination with AP1 it promotes activation, but alone it promotes anergy [4]. |
| **IL2** | 1. HGNC:6001<br>2. PMID:23352221 | AP1 & NFAT & NFkB<br>This node represents the cytokine Interleukin-2. It is a marker of T cell activator, and also a mediator of T cells growth and proliferation. |
| **CD69** | 1. HGNC:1694<br>2. PMID:28507790<br>3. PMID:7665567<br>4. PMID:19841192 | NFkB & HIF1A<br>This node represents the early activation marker of T cells CD69. It is a type II C-type lectin involved in lymphocyte migration and cytokine secretion. |
| **QUIESCENCE** | 1. PMID:23601682 | FOXO & !(AP1 \| NFkB \| NFAT \| cMYC)<br>This node represents the phenotype of resting naive T cells. It is defined as a unique configuration of metabolic and signaling pathways; which results in a basal oxidative metabolism supported mainly by the mitochondria. |
| **ACTIVATION** | 1. PMID:19132916<br>2. PMID:23601682 | AP1 & NFAT & NFkB & GLYCOLYSIS & GLUTAMINOLYSIS & !FAO & FAS & PPP & !ATPSyn<br>This node represents the phenotype of activated T cells. It is defined as a unique configuration of metabolic and signaling pathways; which results in Warburg metabolism of proliferative cells supported mainly by the cytosol. |
| **ANERGY** | 1. PMID:26256793<br>2. PMID:24115444<br>3. PMID:19841171<br>4. PMID:19494254 | NFAT & !FOXO & !AP1 & !NFkB<br>This node represents the phenotype of anergic T cells. It is defined as a unique configuration of metabolic and signaling pathways; which results in an incomplete activation and no response. |
| **METABOLIC_ANERGY** | 1. PMID:26256793<br>2. PMID:24115444<br>3. PMID:19841171 | GLYCOLYSIS & KrebsCycle & !(FAS & GLUTAMINOLYSIS)<br>This node represents the phenotype of metabolically anergic T cells. It is defined as a unique configuration of metabolic and signaling pathways; which results in a cell not ready for proliferation upon stimulation. |
